# Discovery of anti-infective compounds against *Mycobacterium marinum* after biotransformation of simple natural stilbene scaffolds by a fungal secretome

**DOI:** 10.1101/2024.04.18.590143

**Authors:** Jahn Nitschke, Robin Huber, Stefania Vossio, Dimitri Moreau, Laurence Marcourt, Katia Gindro, Emerson F. Queiroz, Thierry Soldati, Nabil Hanna

## Abstract

This study evaluated the efficacy of a high-throughput *Dictyostelium discoideum* – *Mycobacterium marinum* Dd-Mm infection system by first benchmarking it against a set of antibiotics and second in screening a library of natural product (NP) derivatives for anti-infective activity against intracellular *Mycobacterium marinum* (Mm). The study observed no activity of pyrazinamide against Mm, consistent with known resistance patterns, and confirmed other antibiotics, such as rifampicin and bedaquiline, with activity below defined antibacterial susceptibility breakpoints. From screening a small library of NP derivatives, *trans*-δ-viniferins emerged as promising anti-infective scaffolds, particularly two compounds which exhibited an anti-infective activity on Mm during infection but not on Mm in broth, **17** with an IC_50_ of 18.1 µM, and **19** with an IC_50_ of 9 µM). Subsequent exploration via halogenation and structure-activity relationship (SAR) studies led to the identification of derivatives with improved selectivity and potency. The observed anti-infective phenotype may involve mechanisms such as blocking mycobacterial virulence factors or boosting host defense. Furthermore, the study highlights the potential of natural product-inspired derivatization approaches for drug discovery and underscores the utility of the Dd-Mm infection system in identifying novel anti-infective compounds.

**IMPORTANCE:** This study underscores the significance of leveraging natural product-inspired approaches and innovative infection models in search for novel anti-infective compounds. By benchmarking and employing high-throughput *Dictyostelium discoideum*-*Mycobacterium marinum* infection system on a small, focused library of natural product derivatives, the study identified *trans*-δ-viniferins as promising anti-infective scaffolds against *Mycobacterium marinum*, opening potential therapeutic avenues for combating tuberculosis. The findings highlight the value of exploring nature-inspired chemistry for drug discovery and addressing global health challenges.

## INTRODUCTION

Tuberculosis, an infectious disease caused by *Mycobacterium tuberculosis* (Mtb), has tragically claimed the lives of nearly 30 million people across the globe over the past decade, as reported by the World Health Organization (WHO) (Bagcchi, 2023). The tuberculous mycobacteria within the Mtb complex pose a difficult challenge for drug targeting, as these intracellular pathogens reside inside a vacuole in macrophages. Adding to this complexity, the intricate and waxy mycobacteria cell wall serves as an impermeable barrier to effective antibiotic treatment (Alderwick et al., 2015). The quest for new anti-tubercular drugs is an enduring and demanding journey further complicated by Mtb’s ability to enter a metabolically dormant state. Indeed, latent infections are often accompanied by non-heritable antibiotic tolerance, hindering the efficacy of conventional antibiotics and presenting a complicated challenge in disease control (Lewis, 2007). Adding to the gravity of the situation is the swift emergence of Multi-Drug-Resistant (MDR) and Extensively-Drug-Resistant strains (XDR) strains, which demand prolonged antibiotic treatment associated with severe side effects, high costs, and a limited chance of cure (Dheda & Migliori, 2012). Regrettably, despite the escalating threat of antibiotic resistance, the development of new antibiotics has dwindled.

In the pursuit of new anti-Mtb compounds, two primary approaches have been embraced, targeted drug discovery and whole-cell screening. While the main drawback of targeted drug discovery is that the selected target may not be essential *in vivo*, phenotypic screenings have exhibited more promise, giving rise to compounds such as SQ109 which is in clinical trial phase II, (Reddy et al., 2010), bedaquiline (Andries et al., 2005) or delamanid (Matsumoto et al., 2006). Phenotypic screens in whole-cell assays offer numerous advantages over target-based drug discovery, including the identification of novel targets, probing multiple targets simultaneously or detection and exclusion of compounds with low membrane permeability. The fact that 40 years passed before bedaquiline was discovered as a new first-in-class drug (Mahajan, 2013), emphasizes the need for new targets and the importance of phenotypic assays.

Even though SQ109, bedaquiline and delamanid were discovered by phenotypic whole-cell screens on mycobacteria in broth, *in vitro* screening neglects that Mtb is an intracellular, vacuolar or cytosolic pathogen. Consequently, screening approaches should better take into account Mtb’s infection biology by making use of dedicated host-pathogen systems. Recent efforts shifted their focus towards a variety of *in cellulo* infection models, encompassing both cells of animal and non-animal origin. The experimentally versatile Amoebae – *Mycobacterium marinum* (Mm) infection models provide robust and ethically compliant platforms for investigating mycobacterial pathogenicity. Crucially, both Mtb and Mm employ highly similar strategies to establish an infection by hindering phagosome maturation and avoiding killing in macrophages and amoebae, while Mm offers experimental advantages, such as faster growth and being an organism requiring biosafety level 2 infrastructure (Ramakrishnan, 2004; Tobin & Ramakrishnan, 2008). On the host side, two main amoebae systems, *Acanthamoeba castellanii* (E. A. Diop et al., 2018; E. H. A. Diop et al., 2019; Kicka et al., 2014) and *Dictyostelium discoideum* (Dd), are commonly used to study the infectious process caused by various pathogenic bacteria, including Salmonella, Mycobacteria, Legionella, or Pseudomonas (Dunn et al., 2018; Hagedorn & Soldati, 2007; Solomon et al., 2003; Steinert & Heuner, 2005). Both amoebae are professional phagocytes employing evolutionary conserved strategies to capture and kill bacteria. After uptake of Mtb or Mm in macrophages or amoeba, the phagosome becomes a Mycobacteria-Containing Vacuole (MCV), an active interface between the host cell and the mycobacteria, being at the crossroads of host machineries such as membrane damage repair, autophagy, oxidative burst and lysosomal fusion, and virulence factors of the pathogen, such as type VII secretion systems and their effectors. Eventually, the MCV loses its integrity, allowing Mtb or Mm to access the host cytosol (Cardenal-Muñoz et al., 2017; Dunn et al., 2018; López-Jiménez et al., 2018). Amoebae systems and especially Dd offer experimental advantages over animal derived cellular models mainly due to low maintenance cost and ease of genetic manipulation, rendering them ideal platforms for high-throughput approaches. The 3R principles were coined in the 1950s and stand for Replacement, Reduction and Refinement (Russell & Burch, 1959) of animal experiments and the Dd-Mm system is a powerful 3R model system.

Direct screening inside an infected host allows to rapidly exclude compounds that are toxic to the host or have poor pharmacokinetics, thereby significantly contributing to decrease the usually high attrition rate in Mtb drug screens (Huszár et al., 2020). Most importantly, such systems open the possibility to identify host-targeting compounds that boost cell-autonomous defence and bacterial killing. This strategy termed adjunctive or host-directed therapy (HDT) is of particular interest as a strategy against MDR-Mtb (Gries et al., 2020). For example bedaquiline has been shown to activate macrophages in addition to its antibiotic effect (Giraud-Gatineau et al., 2020). The drug’s concentration in lipid droplets of the host cell (Greenwood et al., 2019) might play a substantial role in its mode of action and emphasizes the need of considering host biology when searching for new drugs. Besides unlocking host targets, host-pathogen screening systems also have the potential to guide the way to targets within bacterial effectors, which are specific for the intracellular life of mycobacteria (Gries et al., 2020). Compounds with such an activity have been identified by investigating hits from a host-pathogen screen, which were inactive in a screen on mycobacteria (Rybniker et al., 2014).

Even with the amoeba-Mm infection model as a valuable screening tool, the choice of chemical space to screen is crucial to success. In this regard, natural products (NPs) and their derivatives have played a significant role throughout history in advancing drug therapy, with notable contributions in fields such as cancer and infectious diseases (Newman & Cragg, 2016, 2020). The hegemony of NPs in drug discovery was severely challenged in the 1990s, with a major shift towards synthetic and combinatorial chemistry. However, this shift did not deliver the expected success (David et al., 2015). Recent advances in “omics” techniques, computational methods, and innovative screening models have revitalized the field of phenotypic screens in complex systems, and opened up new avenues of exploration (Rossiter et al., 2017; Wolfender et al., 2019). Despite this advance, several challenges in NP drug discovery remain, such as access to novel biological material, intraspecies variability and supply (David, 2018; Kingston, 2011). An alternative approach that takes advantage of the aforementioned advance is the generation of new NP-like compounds based on NP scaffolds. Such a strategy can take advantage of the use of enzymes to generate derivatives of known NPs that are not or not yet found in nature.

Based on this strategy, we recently presented the use of an enriched enzyme fraction secreted by the fungus *Botrytis cinerea* (“enzymatic secretome”) to transform NPs, such as stilbenes, into more complex structures with improved biological activities against fungi, breast cancer, bacteria and viruses (Gindro et al., 2017; Huber, Koval, et al., 2022; Righi et al., 2020). This approach yielded more than 70 compounds distributed in six scaffolds. Since stilbene derivatives have been reported to have potential anti-TB activity in various *in vitro* assays (Peraman et al., 2021; Reinheimer et al., 2018; Suarez et al., 2017), this library was subjected to an anti-infective screening workflow in the Dd-Mm infection model. The goal of our proof-of-concept study was to develop and benchmark the Dd-Mm system to high-throughput capacity in order to screen a focused library of natural products derivatives for anti-infective and antivirulence activities.

## MATERIALS AND METHODS

### Generation of a stilbene dimer library by chemoenzymatic synthesis

The initial screening library was generated by chemoenzymatic synthesis for a previous study on Wnt pathway inhibition (Huber, Koval, et al., 2022). Briefly, compounds **1**-**4** (99% purity) were purchased from Biopurify Phytochemicals Ltd (Chengdu, Sichuan, China), and dimers **5**-**54** were obtained by enzymatic radical dimerization of **1** and **2** using the enzymatic secretome of *Botrytis cinerea* Pers. (Gindro et al., 2017). Each compound was isolated by semi-preparative high-resolution liquid chromatography (semi-preparative-HPLC) and characterized by nuclear magnetic resonance (NMR), high-resolution mass spectrometry (HRMS) and ultraviolet (UV) spectroscopy. Details on the synthesis, isolation and characterization of the compounds can be found in their original reference (Huber, Koval, et al., 2022).

### Isolation of *trans*-δ-viniferin enantiomers

Both enantiomers of the *trans*-δ-viniferin derivatives **11**, **17**, **19** and **21** were previously isolated for their antibacterial evaluation against *Staphylococcus aureus* (Huber, Marcourt, et al., 2022). This was achieved by using a chiral stationary phase in HPLC. Each enantiomer was characterized by electronic circular dichroism (ECD) and optical rotation to assess its absolute configuration. Details of the isolation and characterization of these enantiomers can be found in the original reference (Huber, Marcourt, et al., 2022).

### Generation of *trans*-δ-viniferin derivatives

The collection of *trans*-δ-viniferin derivatives **55**-**94** was generated for a previous study on antibacterial activity against *S. aureus* (Huber, Marcourt, et al., 2022). Several strategies were employed, including (1) light-induced double-bond isomerization, (2) O-methylation of the phenolic moieties, (3) oxidation of the dihydrobenzofuran ring to benzofuran, (4) radical dimerization of other stilbene monomers and (5) oxidative halogenation. Each compound was isolated by semi-preparative HPLC and characterized by NMR, HRMS and UV spectroscopy. Details of the synthesis, isolation and characterization of the compounds can be found in their original reference (Huber, 2023).

### Antibacterial assay

*M. marinum* WT expressing the *lux* operon (*luxCDABE*) under the *hsp60* promoter (Andreu et al., 2010; Arafah et al., 2013) was cultivated 24 hpi prior to the experiment in a 10 ml volume of 7H9 broth (Becton Dickinson, Difco Middlebrook 7H9) supplemented with 10% OADC (Becton Dickinson) and 0.05% tyloxapol (Sigma Aldrich) and 50 µg/mL kanamycin at 32°C overnight with continuous shaking. The *M. marinum* culture was centrifuged at 300 RPM for 1 minute to sediment large bacterial clumps. The OD_600_ of the supernatant was then measured to determine and subsequently adjust the bacterial density to 3.75*10^5^ bacteria per ml in 7H9 medium. Each well of a 384-well plate (Interchim FP-BA8240) was filled with 20 μL of the bacterial suspension (7.5*10^3^ bacteria), and 0.2 μL of the dissolved compounds were added in a 1:100 dilution to each well to a final vehicle concentration of 0.5% ethanol. Each compound was tested in technical duplicates, and at least 24 technical replicates of control conditions were included: a vehicle control (VC) with a final concentration of 0.5% ethanol and rifabutin at a final concentration of 10 μM as a positive control. The plates were sealed with a gas-impermeable membrane (H769.1, Carl Roth), briefly centrifuged and bacterial growth was monitored using an Agilent BioTek H1 plate reader, an Agilent BioTek BioStack plate stacker, and luminescence was recorded over 72 hours at 32°C with readings taken every hour.

### Anti-infective assay

*D. discoideum* Ax2(ka) expressing mCherry at the *act5* locus (Paschke et al., 2019) was cultured in 10 cm culture dishes (Falcon) in HL5-C medium (Formedium). *M. marinum* M strain expressing the bacterial *lux* operon (*luxCDABE*) was cultured at 32°C under shaking conditions in 7H9 broth (Becton Dickinson, Difco Middlebrook 7H9) containing 0.2% glycerol (Sigma Aldrich), 10% OADC (Becton Dickinson) and 0.05% tyloxapol (Sigma Aldrich). The day before the experiment 10^7^ amoebae were plated in a 10 cm Petri dish. On the day of the experiment, infection was performed at MOI 25 by spinoculation in HL5-C and subsequently washing off extracellular bacteria (Mottet et al., 2021). Then, the cell suspension of infected amoebae was detached from the culture dish and resuspended in HL5-C with 5 U/mL penicillin and 5 μg/mL of streptomycin (Gibco) to inhibit extracellular growth of bacteria. The density of infected cells was quantified using a Countess (Thermo Fisher Scientific), and 1×10^4^ cells in 20 µL of HL5-C with penicillin and streptomycin were seeded into each well of a 384-well plate (Interchim FP-BA8240). Test compounds resuspended in 50% ethanol were tested in technical triplicates and added in a 1:100 dilution to a final vehicle control concentration of 0.5% ethanol. The plates were sealed (H769.1, Carl Roth), briefly centrifuged and fluorescence and luminescence were monitored using an Agilent BioTek H1 plate reader and an Agilent BioTek BioStack plate stacker in a temperature-controlled environment (set to 24°C) over 72 hours with readings taken every hour.

For both assays, growth curves were obtained by measuring the luminescence and fluorescence as a proxy for bacterial growth and host growth, respectively, for 72 hours with time-points taken every hour. The “normalized residual growth” was computed by calculating the area under the curve (AUC, trapezoid method) and normalization to the vehicle control (0.5% ethanol, set at 1) and a baseline curve (set at 0). The baseline curve was calculated by taking the median of the first measurement of all wells in a plate and extrapolating it over the full time course. The threshold for hit detection was arbitrarily fixed at a cut-off of normalized residual growth < 0.5. Normalized values were averaged over technical and biological replicates (all experiments at least n = 3 and N = 3) before estimating the IC_50_. For IC_50_ estimation, a robust 4PL regression was used, constraining the top value to 1 and, if necessary, the bottom value to −2. This was performed using GraphPad Prism (Version 8.0.1). The MIC was determined as the lowest experimental concentration, where the averaged normalized residual growth plus one standard deviation included 0. For every assay plate the robust Z’-factor was calculated using the vehicle control and the positive control.

### Anti-infective assay with High-content microscopy

The procedure described here was optimized from (Mottet et al., 2021). To monitor both the host and pathogen during the course of infection in a quantitative manner, *D. discoideum* Ax2(ka) cells expressing mCherry knocked-in at the *act5* locus (Paschke et al., 2019) were infected as described above with *M. marinum* WT stably expressing GFP (Msp:12-GFP). As described above, the day before the experiment, 10^7^ amoebae were plated in a 10 cm Petri dish. On the day of the experiment, infection was performed, as described above, in HL5-C at MOI 25 by spinoculation. This was followed by eight washes in HL5-C to clear a maximum of non-internalized bacteria. The infected amoebae were detached mechanically and resuspended in HL5-C with 0.75 U/mL penicillin and 0.75 μg/mL of streptomycin (Gibco). Compounds were diluted in HL5-C to a 10x stock of the final assay concentration and pre-plated in 20 µL into the respective wells of a 96-well μ-plate (iBIDI 96-well μ-plate, 89626). Subsequently, 7500 cells per well were added in a volume of 180 µL. Every condition has been plated in at least three technical replicates and at least three biological replicates. The 96-well plate was sealed with a gas permeable seal (4titude, 4ti-0516/96) and centrifuged briefly.

Microscopy images were acquired at 25°C with a 40X plan Apo objective at selected time points over 48 h on the ImageXpress Micro C® automated microscope (Molecular Devices) in wide field mode. Focus was obtained automatically by autofocusing on the well bottom and plate bottom, at every well and every time point. In a grid of 4×5, 20 images per well, corresponding to a range of 500-6000 cells, have been taken using the Texas-Red channel for *D. discoideum* expressing mCherry and FITC channel for *M. marinum* expressing GFP in each experiment. Image analysis was performed using MetaXpress Custom Module editor software. Objects were effectively segmented by size and intensity. More precisely, amoebae were segmented using a “find blob module” using the Texas-Red channel. To segment the bacteria, the FITC image was first subjected to a light deconvolution with a top hat filter and then segmented using a “find blob module”. Intracellular bacteria and infected amoebae were determined by associating the mask of the amoebae and the bacteria. Several parameters such as intracellular and extracellular bacterial area sum, amoebae area sum, averaged and integrated fluorescence intensity and feature counts of bacteria and amoeba were extracted. The growth curves for the area sum of uninfected amoebae, infected amoebae and intracellular bacteria of independent biological replicates were averaged and plotted using GraphPad Prism (Version 8.0.1).

## RESULTS

### Benchmarking the Dd-Mm high-throughput system

The high-throughput *Dictyostelium discoideum - Mycobacterium marinum* (Dd-Mm) infection model was benchmarked by monitoring dose-response curves (DRCs) for seven antibiotics used in TB chemotherapy (Fig. 1). First, we used an anti-infective assay, i.e. Mm-infected Dd, and second an antibacterial assay, i.e. Mm growing in broth. We used a Dd strain expressing mCherry to measure the growth of the amoebae in parallel with the intracellular growth of a bioluminescent Mm strain. In both assays, the luminescence serves as a proxy for Mm mass and metabolic activity. The bacteria or infected amoebae were plated in 384-well plates, treated with descending concentrations of each antibiotic and monitored for 72 hours using a plate reader. The average robust Z’-factor of the respective benchmarking experiments was 0.73 for Mm in infection and 0.61 for Mm in broth. The antibiotics rifampicin, isoniazid, ethambutol, bedaquiline, ethionamide and rifabutin exhibited sterilizing activity in both assays at their respective maximal tested concentration (Fig. 1A-I). In addition, we observed a slight pro-infective effect of isoniazid between 0.4 and 3.3 µM (Fig. 1E) and also of pyrazinamide, while the latter showed otherwise no activity against Mm (Fig. J). Moreover, rifampicin and ethambutol showed dramatically different activities in infection and in broth with MIC of 10 and 1.1 µM, respectively for rifampicin (Fig. 1D), and MIC of 30 and 3.3 µM, respectively for ethambutol (Fig. 1F). Rifabutin and ethionamide also exhibited moderate differences of activity on Mm in infection and Mm in broth. Both, rifampicin and rifabutin affected amoebae growth slightly at the highest tested concentrations, but, as illustrated for rifampicin in panel C, did not prohibit Dd from reaching a plateau in its growth phase.

**Fig. 1.**
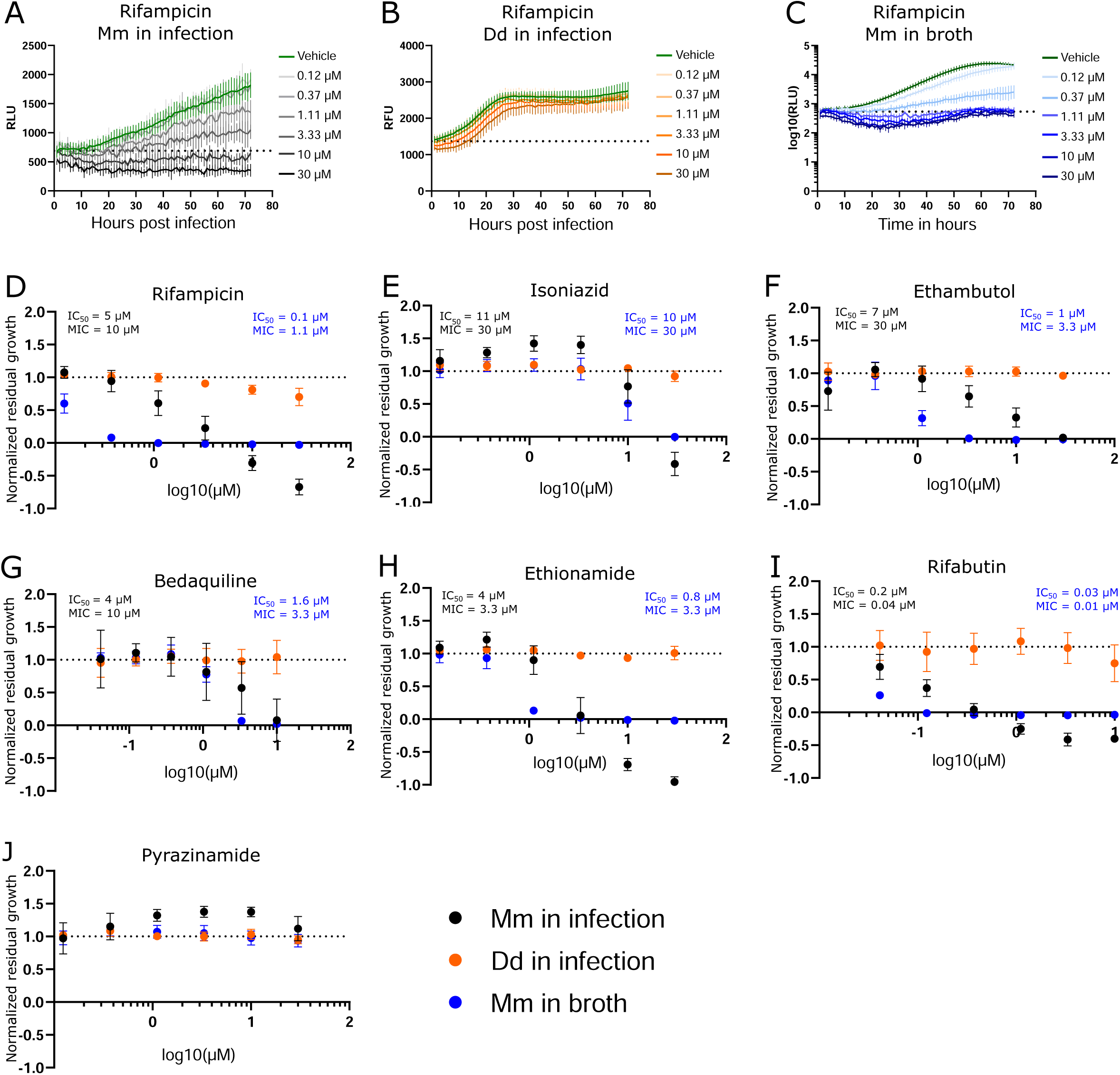
Dose response curves of benchmarked antibiotics. A selection of antibiotics used for treatment of a TB infection were benchmarked in the Dd-Mm high-throughput system, in infection and in broth. In panels **A**, **B** and **C**, 72 hours growth curves of Mm in infection, Dd in infection and Mm in broth, respectively, at different concentrations of rifampicin are shown. The dashed line represents the median over the first measurements of all wells of the respective experiment. On the y-axis are random luminescence units (RLU) or random fluorescence units (RFU). The former were log10 transformed in panel **C**. The corresponding dose-response curve of Rifampicin is shown in panel **D**. The dose-response curves depicted in panel **E** (isoniazid), **F** (ethambutol), **G** (bedaquiline), **H** (ethionamide), **I** (rifabutin) and **J** (pyrazinamide) were created from growth curves in analogous manner. Black points show Mm in infection, orange points Dd in infection and blue points Mm in broth. Log10 of the test concentration in µM is shown on the x-axis. A dashed line at y = 1 represents the normalized residual growth of the vehicle control. Depicted are means of at least three biological replicates and the respective standard deviations. Black and blue text inlays show the MIC and IC_50_ of the anti-infective or the antibacterial assay, respectively.

### Primary screening of the stilbene monomer and dimer library in a high-throughput and dual readout growth inhibition assay

Next, we selected a library of compounds obtained through chemoenzymatic dimerization allowing structure activity relationship (SAR) investigations, while being relatively diverse. The compound structures can be classified into five scaffolds: *tran*s-δ-viniferin, pallidol, leachianol, restrytisol and acyclic dimer. Each scaffold is derived from a different coupling reaction, as described in our previous article (Huber, Koval, et al., 2022). Restrytisol, leachianol and acyclic dimer scaffolds are further classified according to their relative stereochemistry. Two parameters explain the diversity of compounds obtained in each scaffold: First, the starting materials: resveratrol, pterostilbene, or a mixture of the two, which lead to hydroxylated and *O*-methylated derivatives, and second, the cosolvent. The leachianol and acyclic dimer scaffolds are the only ones capable of incorporating a solvent molecule during their synthesis by a solvolysis reaction, i.e., a nucleophilic attack of the alcohol on an sp^2^ carbon of an intermediate. This explains the large number of members in these two series. In addition to these 50 stilbene dimers, four stilbene monomers, which were used in generating dimers or derivatives of the initial library (including resveratrol, pterostilbene, pinostilbene and isorhapontigenin) were included in the library (Fig. 2A).

**Fig. 2.**
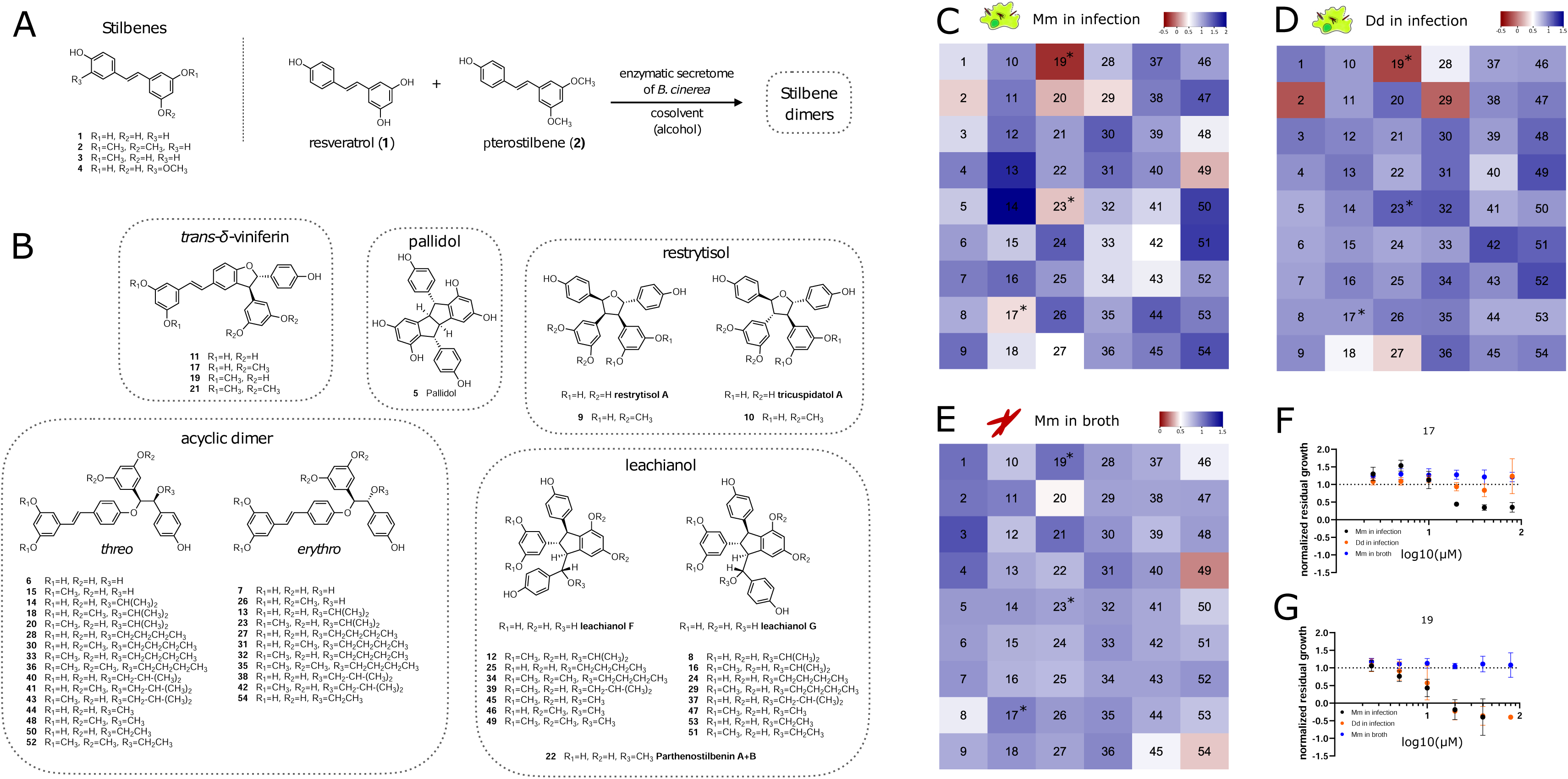
Overview of compound structures in the library and results of the screening. **A**: Stilbene monomers (**1-4**) and biotransformation reaction with the enzymatic secretome of *B. cinerea*. **B**: Structures of the stilbene dimers obtained by biotransformation of resveratrol and pterostilbene (**5-54**). Heatmaps in **C**, **D** and **E** colour code normalized residual growth of Mm in infection (**C**), Dd in infection (**D**) and Mm in broth (**E**) under treatment with compounds at 20 µM each. Values below the cut-off of 0.5 are displayed in shades of red, while values above the cut-off are displayed in shades of blue. Numbers in the heatmap correspond to the compounds described in panels **A** and **B.** Compounds meeting the cut-off at Mm in infection, but not in broth (i.e. Mm in infection ≤ 0.5 and Mm in broth ≥ 0.5) after a validation in a dose-response curve are marked with an asterisk in all three heatmaps. **F** and **G** depict these dose-response curves of **17** and **19**, respectively, of which both are analogs of the same scaffold, the *trans*-δ-viniferins. The corresponding dose-response curve for **23** can be found in Fig S1. The log10 of the concentration in µM is shown on the x-axis; a dashed line at y = 1 represents the normalized residual growth of the vehicle control. Black points show Mm in infection, orange points Dd in infection and blue points Mm in broth. Shown are means of at least three biological replicates and the respective standard deviations. Note that in **G**, for visual clarity the highest dosage (80 µM, normalized residual growth of −2.2) was omitted for Mm in infection.

Screening on infected cells is more stringent than screening on Mm in broth, since compounds with poor pharmacokinetics are excluded. By utilizing the Mm in broth screen as a counter screen instead of a validation, we additionally focused on compounds which act specifically during infection. We subjected the library to the anti-infective and antibacterial high-throughput assays, analogously to the procedure applied in Fig. 1. The hit cut-off for any readout was set at 50% of residual growth. This allowed a hit classification for each compound in the library and for each readout: Mm in infection, Dd in infection and Mm in broth. The average robust Z’-factors for this screen were 0.71 for Mm in infection and 0.74 for Mm in broth.

One of the four monomers **1-4**, pterostilbene (**2**) showed anti-infective activity (0.28 of normalized residual growth in Mm in infection), but inactive on Mm in broth (Fig. S1B). However, this compound exhibited a strong Dd growth inhibition (−0.16 of normalized residual growth of Dd in infection), which is a possible indirect cause for its activity on Mm. Pterostilbene was tested in a DRC, revealing that activity on Dd in infection occurred already at lower concentrations than activity on Mm in infection (IC_50_ estimate of 11.1 µM and 19.8 µM respectively, Fig. S1B, Table S1), weakening a possible causal link between activity on Dd and Mm during infection. The other monomers were also tested in a DRC, confirming the lack of activity of resveratrol (**1**, Fig. S1A) and isorhapontigenin (**4**, Fig. S1D) even at high concentrations, while pinostilbene (**3**, Fig. S1C) was active on Mm in infection, at high concentrations (> 20 µM).

Several stilbene dimers were hits on Mm in infection, in particular **17** (with a normalized residual growth of 0.34), **19** (−0.32), **20** (0.20), **23** (0.23), **27** (0.47), **29** (0.35), **42** (0.47) and **49** (0.19) (Fig. 2C, Table S1). In broth, three of these dimers had an antibacterial activity with normalized residual growth below 0.5: **20** (0.47), **49** (0.27) and **54** (0.38) (Fig. 2E), whereas **17**, **19**, **23**, **27**, **29** and **42** were shown to be anti-infectives without antibiotic activity. Among these six compounds, **19** (normalized residual growth of −0.22), **27** (0.32) and **29** (−0.10) were also inhibitors of Dd growth (Fig. 2D, Table S1). The other three dimers, **17** (normalized residual growth of 0.91), **23** (1.1) and **42** (1.2) were inactive on Dd (Fig. 2D, Table S1).

All six dimers were subsequently validated in the anti-infective assay in a DRC. All of them exhibited a dose-dependent activity, but only **17**, **19** and **23** passed again the 0.5 cut-off for Mm in infection at 20 µM (Fig. 2FG and Fig. S1E). A high anti-infective activity was confirmed for **19** (IC_50_ = 9 µM, Table S1), but was seen as potentially correlated to a high activity on Dd growth (IC_50_ = 12.1 µM, see Table S1). For **17**, a lower maximal effect on Mm in infection was observed (normalized residual growth of 0.3 for **17** compared to −0.2 for **19**), but also a much lower activity on Dd growth (IC_50_ = 18.1 µM for Mm in infection, an estimate for Dd in infection was not possible due to the shape of the curve, Fig. 2F, Table S1) indicating a better selectivity of **17** towards the bacteria compared to **19**. Both compounds did not show any growth inhibition of Mm in broth. Regarding compound **23**, activity at 20 µM was confirmed by a DRC with an IC_50_ of 10.4 µM and no activity on Dd growth (Fig. S1E).

Surprisingly, few compounds (**13**, **14**, **50**, **51**, and **54)** had a pro-infective activity and enhanced Mm growth during infection with a normalized residual growth value greater than 1.5 (Fig. 2C, Table S1). This feature was exclusively identified during infection and not in broth. In the scope of this study, we focused on conditions that limit Mm growth, therefore, these compounds were not further explored.

Both **17** and **19** belong to the *trans*-δ-viniferin scaffold, while **23** is an acyclic dimer. Besides **17** and **19**, which both bear two *O*-methyl groups in a di-*O*-metyhlated ring, **11** and **21** also belong to the *trans*-δ-viniferin scaffold (Fig. 2B), but with either no (**11**) or four (**21**) *O*-methyl groups, as two di-*O*-metyhlated rings. Both **11** and **21** showed no activity at 20 µM on Mm in infection with normalized residual growth of 1.1 and 1.4, respectively, or in broth both with values of 1. Compounds **13** and **14** are both closely related to the acyclic dimer **23** but have both no *O*-methyl groups and are both inactive in infection (1.7, 2.0 respectively) and in broth (0.85, 0.90 respectively) (Table S1).

In summary, the primary screen revealed **20** and **49** to be active both against extracellular and intracellular bacteria, whereas **2**, **17**, **19** and **23** were validated anti-infectives without antibiotic properties, partly with activity against Dd (**2** and **19**), and **54** was exclusively active against Mm in broth. Pterostilbene (**2**) is a monomer, while **17** and **19** both belong to the *trans*-δ-viniferin scaffold and **23** is an acyclic dimer. Both, *trans*-δ-viniferin and the acyclic dimer families contain very similar but inactive molecules that differ in the number and position of *O*-methyl groups. Based on these different activity patterns of the *trans*-δ-viniferins we subsequently focused on exploring this set of analogs.

### High-content microscopy characterization of two hit compounds

After identifying and validating the *trans*-δ-viniferin derivatives **17** and **19** as anti-infectives without antibiotic activity, they were further validated using time-lapse high-throughput and high-content microscopy. While the plate reader assay provided aggregated data of a partially infected cell population, High-content microscopy allowed us to focus exclusively on infected cells in the population. Both uninfected Dd and Dd infected with Mm were treated with 20, 10 or 2.5 µM of **17** or **19** and the infection was monitored by collecting microscopy images at five time points for up to 48 hours. The top concentration of 20 µM was chosen to be consistent with the primary screen whereas the dilutions were chosen to observe an effect similar to the vehicle control at the lowest dose. Mm and Dd were both segmented (Fig. 3AB) and the area of Dd (Fig. 3CD), infected Dd (i.e. Dd overlapping with a bacterium, Fig. 3EF) and the area of intracellular bacteria, (i.e. Mm overlapping with Dd, Fig. 3GH) were quantified. For all quantified parameters, a dose-dependency was observed for **17** and **19**. The growth of uninfected Dd was more affected by **19** than by **17** (Fig. 3CD), while the growth of infected Dd (Fig. 3EF) and intracellular Mm (Fig. 3GH) showed the same pattern. Overall, the growth of intracellular Mm and infected Dd was almost completely inhibited by **19**, while growth of uninfected Dd was also significantly reduced. For **17**, this effect was weaker and, in addition, the restriction of uninfected Dd was less pronounced than the effect on intracellular Mm and infected Dd. In summary, validation of the activity of **17** and **19** using high-content microscopy confirmed that **17** was more selective for the intracellular pathogen over the host, although the overall activity was lower than that of **19**.

After validating **17** and **19** as anti-infective compounds against Mm, we wondered if an interaction between **17** and established antibiotics could be determined. We tested this compound for synergy with rifampicin and isoniazid in a checkerboard assay and additionally tested for synergy between rifampicin and isoniazid. In brief, we did not find striking synergy but rather the pro-infective effect of **17** at low dosages (see Fig. 2C) dominated the antibiotic effect of both drugs (between 1.25 and 10 µM of **17**, see Fig. S3CE).

### Evaluation of *trans*-δ-viniferin derivatives

After validating the activity of **17** and **19** by high-content microscopy, their structure-activity relationship was further investigated, as **17** and **19** showed different activity and selectivity profiles despite having very similar structures. First, the impact of their absolute configuration was investigated. Indeed, the stilbene dimers (**5-54**) are all obtained as racemic mixtures by phenoxy radical coupling. Since enantiomers often show different activities, the previously separated enantiomers of the four *trans*-δ-viniferin derivatives (Fig. S2A, **11**, **17**, **19**, **21**) (Huber, Marcourt, et al., 2022) were evaluated for their anti-Mm activity during infection. The results, shown in Fig. S2BCD, indicated similar activity for both enantiomers of **17** and **19**. Both enantiomers of **21** remained inactive, while a slight difference was observed between the enantiomers of **11**. The latter result has not been further investigated due to the low level of activity observed.

Next, a series of derivatives of *trans*-δ-viniferin was generated and tested. These derivatives were obtained by light isomerization of the double bond (compounds **55-58**, Fig. 4A), by *O*-methylation of the phenolic moieties (compounds **59-64**, Fig. 4B) and by oxidation to generate benzofuran derivatives (compounds **66-69**, Fig. 4C). A methoxylated *trans*-δ-viniferin derivative (**65**) was also obtained by radical coupling of isorhapontigenin (**4**, Fig. 4D). *Cis* isomerization and oxidation were performed to change the general shape of the molecule, the former reorienting a diphenol ring, and the latter making the structure planar. On the other hand, *O*-methylation allowed to study in detail the influence of the number and position of *O*-methyl groups, whose importance has been highlighted in the primary screen by the difference in activity between compounds **11** (no *O*-methyl group, inactive), **17** and **19** (two *O*-methyl groups, active), and **21** (four *O*-methyl groups, inactive).

Among the derivatives, the two chemical reactions that change the global shape of the molecules were first investigated. In the case of a specific protein target, such a change is expected to strongly influence the activity. The *cis* isomers (Fig. 4A) showed a very close activity to the parent compounds (Mm in infection IC_50_ = 18.9 µM for **17**, 19.7 µM for **56** (*cis* isomer); 9.0 µM for **19**, 9.8 µM for **57** (*cis* isomer)). Similarly, the *cis* isomers **55** and **58** of the inactive *trans*-δ-viniferin **11** and **21**, respectively, were also inactive at 20 µM (Table S1). For the benzofuran derivatives, the change in activity was again weak for **17** (19.6 µM for **67,** Fig. 4G, Table S1), but a decrease in activity was observed for **19** (20.1 µM for **68,** Table S1, Fig. 4H). The benzofuran derivative of **21** (**69**) remained inactive (Fig. S1G, Table S1), while the corresponding derivative of **11** (**66**) showed nonspecific activity at higher dose than **17** and **19** (Mm in infection IC_50_ estimates of 29.7 µM, Dd in infection IC_50_ estimates of 26.2 µM; normalized residual growth of Mm in infection at 20 µM of 0.55) (Fig. S1H). A direct comparison with the DRC of **11** was not made because **11** was considered inactive at 20 µM in the primary screen (normalized residual growth of Mm in infection at 20 µM of 1.08). The complete IC_50_ data are given in Table S1.

Already the data from the primary screen showed that the number and position of *O*-methyl groups was important for the activity and selectivity of the compounds. Indeed, **17** and **19** which have both two *O*-methyl groups, showed different activity and selectivity in our assay. This could be due to the distinct positions of these groups. The impact of *O*-methyl groups was further supported by the decreased anti-infective activities of **11** (no *O*-methyl group) and **21** (four *O*-methyl groups). The importance of the *O*-methyl groups was therefore systematically investigated. Mono-*O*-methylated derivatives (**59-61**) were found to have weaker activities (less than 50% growth reduction of Mm in infection at 20 µM) and were therefore not investigated further (Table S1).

On the other hand, the di-*O*-methylated derivatives **62-64** showed an improved activity on Mm in infection (Fig. 4IJK) compared to **17** and **19**. Compound **62**, showed a stronger maximal effect than **17** (−0.6 and 0.4, respectively), and its IC_50_ was slightly improved compared to **17** (13.4 µM and 18.9 µM, respectively, Fig. 4I). Also **63** and **64** showed an improved maximal effect (−0.4 for **63** and 0 for **64**), and the IC_50_ improved by about a factor of two, compared to **17** (8.6 and 8.0 µM for **63** and **64**, respectively, Table S1). Moreover, **62**, **63** and **64** showed a reduced activity on the host during infection compared to **19**, while maintaining a similar effect on Mm in infection (lowest measured value of Dd in infection 0.42, 0.8 and 0.62, respectively; −0.40 for **19**). Notably, none of the modifications induced a gain of activity against Mm in broth, confirming the anti-infective property of the scaffold. On the other hand, compound **65**, corresponding to a dimethoxylated derivative of **11** (keeping the free phenol groups) did not reduce Mm growth during infection, but rather showed pro-infective activity at 20 µM and was therefore disregarded (Fig. S1).

After identifying **62**, **63** and **64** as *trans*-δ-viniferin derivatives with superior potency and selectivity over the initial screening hits **17** and **19**, the activity profiles of the two most selective compounds **62** and **63** were validated using high-content microscopy (Fig. 5). Similar to Fig. 3, uninfected and infected Dd were treated with 20, 10 and 2.5 µM of **63** or **64** and cells were monitored for 48 hours. Uninfected Dd, infected Dd and intracellular Mm were segmented (Fig. 5AB) treated with compounds and/or the vehicle. Quantification of uninfected Dd confirmed only a slight growth inhibition at 20 µM for **63** and **64** (Fig. 5 CD), whereas the infection was well controlled by treatment with each of the compounds (Fig. 5 EFGH).

**Fig. 3.**
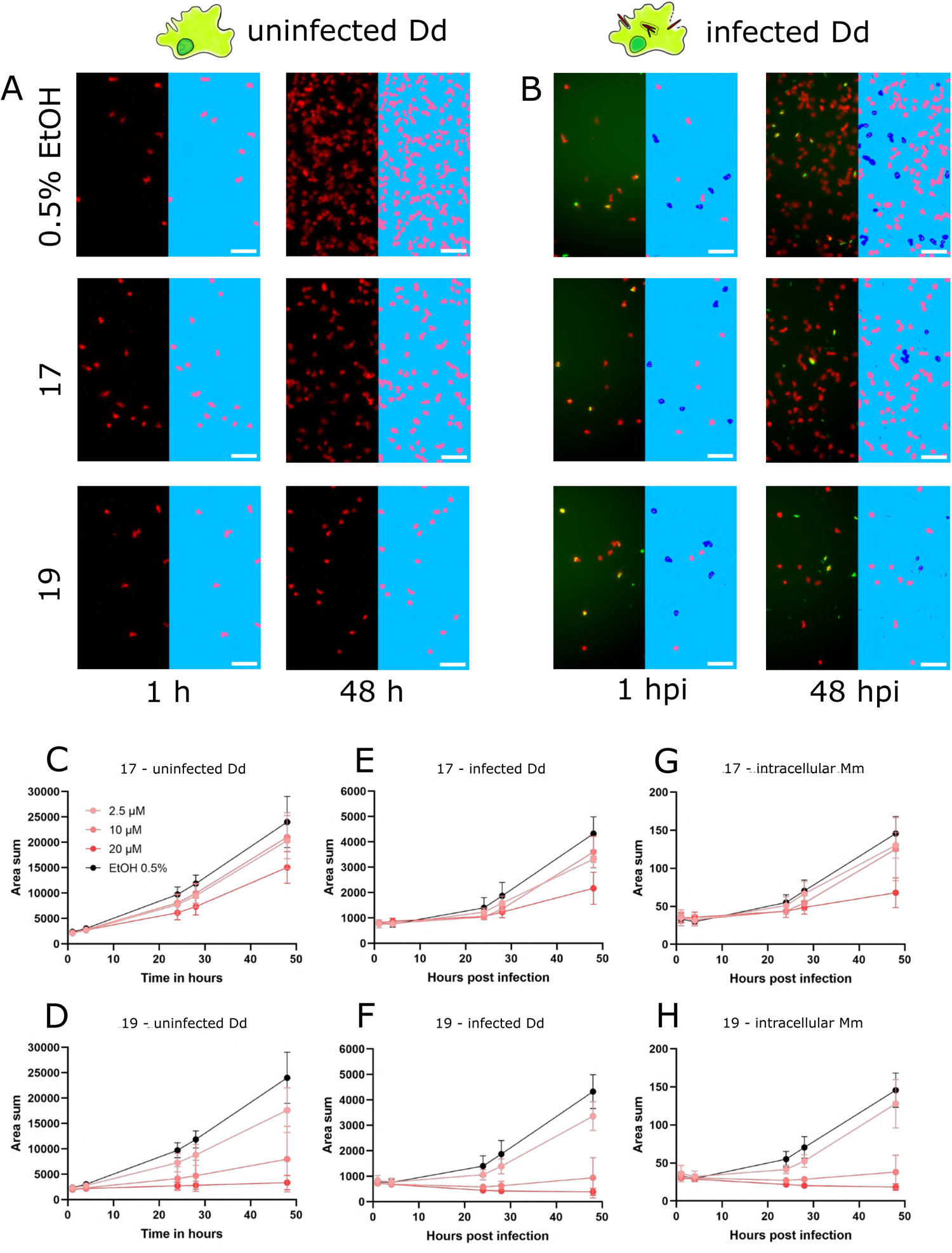
Characterization of primary hits with high-content microscopy. Panels **A** and **B** show high-content microscopy panels illustrating the effects in infected and uninfected amoebae under treatment with **17** and **19** at 20 µM or 0.5% ethanol as the vehicle control at 1 and 48 hpi. The left parts of the panels show the microscopy image, while the right parts of the panels show the corresponding segmentation. Uninfected Dd are shown in pink, infected Dd are shown in dark blue, intracellular bacteria are shown in white and extracellular bacteria are shown in black. Quantifications over the full time course and at concentrations of 5, 10 and 20 µM of **17** and **19** are depicted in **C** to **H**. **C** and **D** show the segmented area of uninfected Dd over time. **E** and **F** show the segmented area of Dd determined to be infected by Dd-Mm area overlap, over time. **G** and **H** show the segmented area of Mm determined to be intracellular by Dd-Mm area overlap, over time. Data points represent the average of three independent biological replicates, error bars are SDs.

**Fig. 4.**
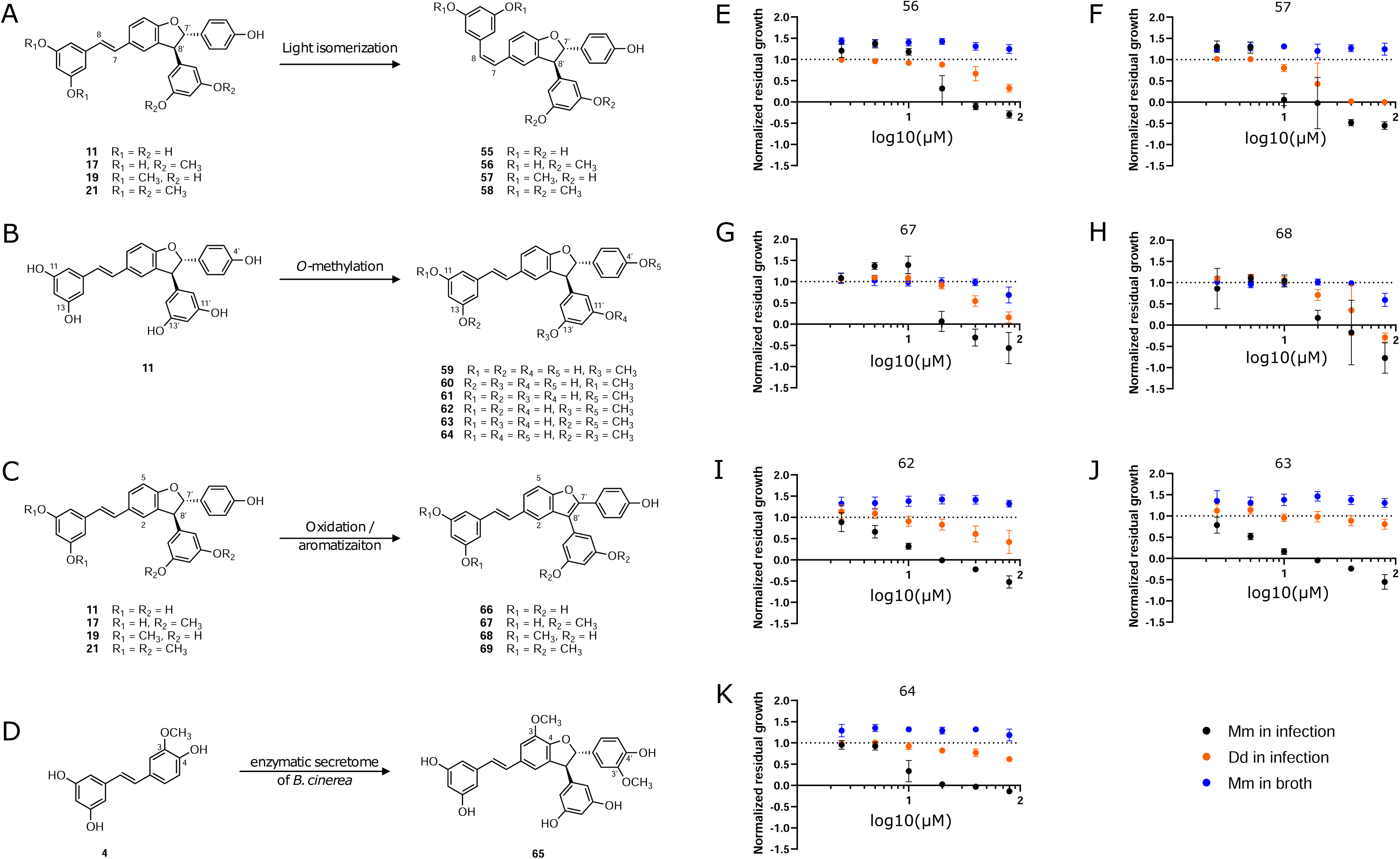
Synthesis of *trans*-δ-viniferin derivatives and corresponding structure activity relationship. Panel **A** shows light-isomerization leading to cis isomers **55-58**. **B** shows *O*-methylation of **11** leading to mono-*O*-methylated derivatives **59-61** and di-O-methylated derivatives **62-64**. **C** shows oxidation resulting in planar structures with a benzofuran moiety, **66-69**. **D** shows a radical coupling reaction of isorhapontigenin (**4**) leading to a dimer with two added methoxy groups (**65**) compared to trans-δ-viniferin (**11**). Panels **E-K** show dose-response curves of normalized residual growth of the aforementioned derivatives. **E** and **F** show *cis* isomers **56** and **57**, **G** and **H** benzofuran derivatives of **17** and **19**, **67** and **68**, respectively. **I**, **J** and **K** show di-*O*-methylated derivatives **62**, **63** and **64**. Black points show Mm in infection, orange points Dd in infection and blue points Mm in broth; Log10 of the concentration in µM is shown on the x-axis. A dashed line at y = 1 represents the normalized residual growth of the vehicle control. Depicted are means of at least three biological replicates and the respective SDs.

**Fig. 5.**
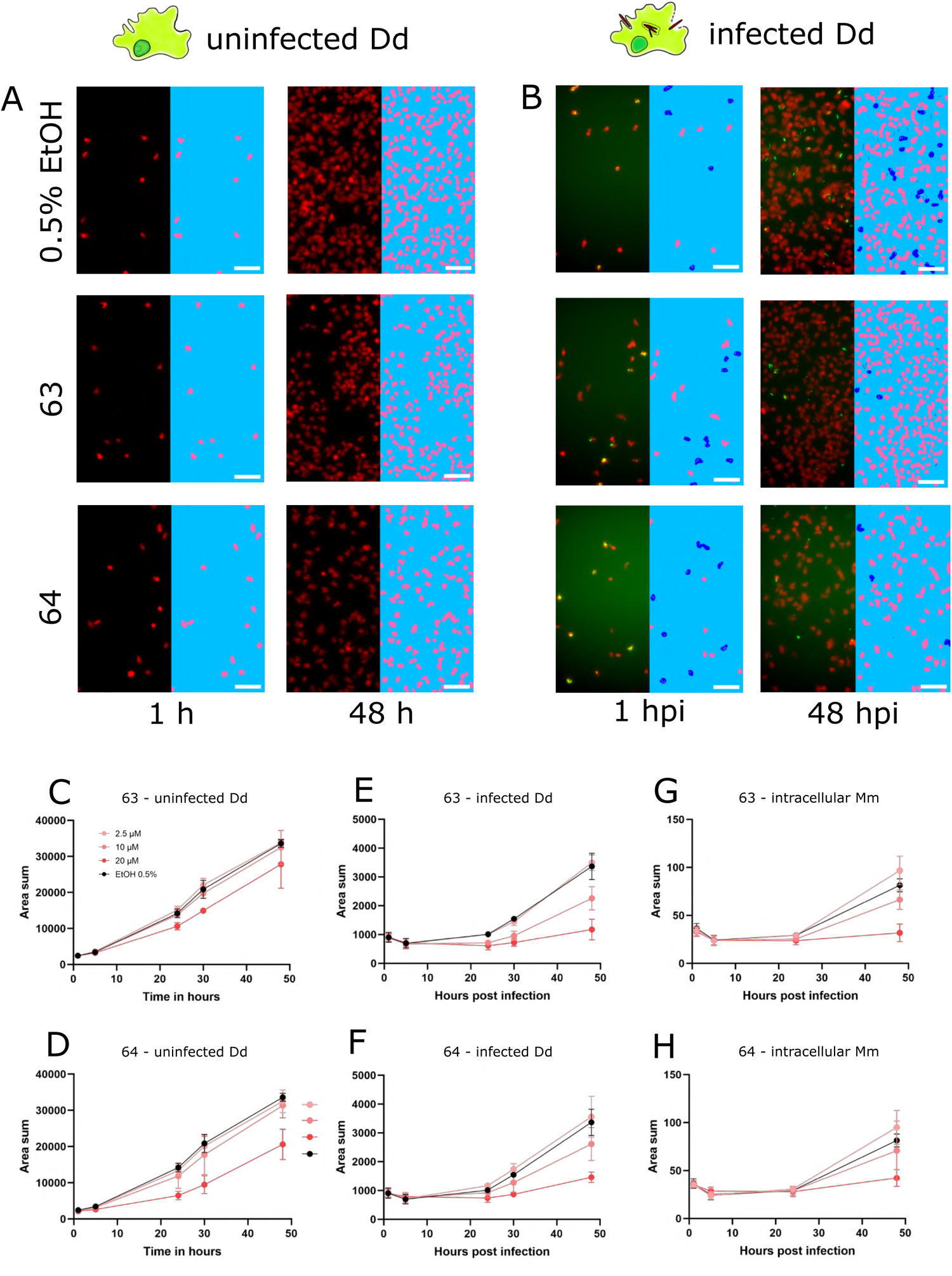
Characterization of improved derivatives with high-content microscopy. Panels **A** and **B** show panels from high-content microscopy, illustrating effects in infected and uninfected amoebae under treatment with **63** and **64** at 20 µM or 0.5% ethanol as the vehicle control at 1 and 48 hpi. The left part of the panels shows the microscopy image, while the right part of the panels shows the segmented cells. Uninfected Dd are shown in pink, infected Dd are shown in dark blue, intracellular bacteria are shown in white and extracellular bacteria are shown in black. Quantifications over the full time-course at concentrations of 5, 10 and 20 µM of **63** and **64** are depicted in **C** to **H**. **C** and **D** show the segmented area of uninfected Dd over time. **E** and **F** show the segmented area of Dd determined to be infected by Dd-Mm area overlap, over time. **G** and **H** show the segmented area of Mm determined to be intracellular by Dd-Mm area overlap, over time. Data points represent the average of three independent biological replicates, error bars are SDs.

Overall, the confirmation of a reduced effect of **63** and **64** on the growth of Dd compared to **19**, while having a similar potency as **19,** and higher than **17**, against Mm in infection render these two derivatives promising hits for further investigation. Interestingly, these four compounds are all *trans*-δ-viniferin isomers with two *O*-methyl groups, the only difference being their positions: both methoxy groups on a single ring for **17** and **19**, or on two different rings for **63** and **64**. The di-*O*-methylated ring connected to the conjugated double bond system in **19** seemed to contribute to activity against Dd, whereas the presence of at least one mono-*O*-methylated ring was beneficial both for maximum effect and selectivity. On the other hand, the other derivatives, *cis* isomers and benzofuran derivatives in particular, did not allow to draw clear conclusions about the structure-activity relationship. The *cis* isomerization did not affect the activity of any of the parent compounds (**11**, **17**, **19** and **21**), while the oxidation to benzofuran resulted in either no or only a small change (**11**, **17**, **21**) or a decrease in activity (**19**).

In addition to modifications on the double bond, the ring system or *O*-methylation, we also halogenated compounds **11**, **17**, **19** and **21** with chloride, bromide and iodine at different positions (Table S1) and subsequently performed preliminary experiments of these derivatives in 96 well plates in infection (n = 3, N = 2) and in broth (n = 3, N = 1). The experiment in broth confirmed no gain of antibiotic activity of any analog (Table S1). The experiments in infection also showed no gain of activity but revealed several inactive analogs. Although no completely consistent pattern emerged, it appeared that halogenation at the diphenol rings had the most impact on activity. In particular double halogenation, e. g. in the case of the inactive analogs of **17**, **77** (di-brominated in positions 10 and 14) and **80** (di-chlorinated in positions 10 and 14) (Table S1), decreases the predicted *p*ka of the hydroxyl groups from approximately 9 to almost 7, rendering them partly deprotonated in the cell. The two inactive analogs of **19**, **83** and **89** also show at least one halogenation at the diphenol ring, whereas halogenation on the di-*O*-methylated ring, such as in **85** and **86** did not decrease activity (Table S1).

## DISCUSSION

In this study, we first benchmarked the high-throughput Dd-Mm infection system and subsequently deployed it in a phenotypic screen on a small library of natural product derivatives. We found *trans*-δ-viniferins to be an anti-infective scaffold, with activity specific to Mm in infection. In particular compounds **17** and **19** stood out, which we subsequently explored by halogenation and further SAR on stereochemistry, planarity and *O*-methylation. Although we did not find striking and consistent patterns in this SAR, we confirmed anti-infective activity and higher potency for three derivatives, **62**, **63** and **64**.

During benchmarking the Dd-Mm system, we observed no activity of pyrazinamide both in infection and in broth (Fig. 1J). This is in accordance with Mm being known to be naturally resistant to pyrazinamide (Aubry et al., 2000; Finkelstein & Oren, 2011). Resistance to isoniazid has also been reported (Aubry et al., 2000; Lorian, 2005), and in both assays isoniazid had the highest observed MIC (30 µM or 4.1 µg/mL). The National Committee for Clinical Laboratory Standards (NCCLS) defined breakpoints of antibacterial susceptibility for MIC_90_ from clinical isolates. Breakpoints for Mm were defined for rifabutin (2 µg/mL), ethambutol (4 µg/mL) (Woods & Clinical and Laboratory Standards Institute, 2011) and ethionamide (5 µg/mL) (Lorian, 2005). MICs observed in this study are all below the respective breakpoints, with the exception of ethambutol in infection (30 µM or 6.1 µg/mL). Indeed, ethambutol and rifampicin showed a large difference of activity between infection and growth in broth. Such a pattern was reported for rifampicin in Raw264.7 (Brodin et al., 2010) and THP-1 macrophages, due to export from the host cell via P-gp, but specifically not for ethambutol (Hartkoorn et al., 2007). Dd possesses an array of ABC transporters and is known for rapid drug export (Anjard et al., 2002; Miranda et al., 2013), emphasizing its stringency as a drug screening model in regard to pharmacokinetics.

After screening a library of stilbene dimers in the Dd-Mm system, we detected mostly anti-infective activities in contrast to classical antibiotic compounds, characterized by no activity in broth throughout the tested concentration range. Subsequently, we investigated the scaffold via targeted SAR and found derivatives with improved selectivity and maximal effect. This demonstrates not only the power of the Dd-Mm system as a 3R infection model, but also the value of exploring natural chemistry by novel approaches of derivatization. The observed anti-infective phenotype might be caused by accumulation inside Dd, which we speculate is unlikely, since Dd is known to rapidly export xenobiotics through its array of ABC transporters (Anjard et al., 2002; Miranda et al., 2013), as it seems to be the case for rifampicin. An anti-infective phenotype is thus likely to be caused by an antivirulence or host-directed defence booster mode of action (Gries et al., 2020).

Previous work has identified host-directed compounds in mammalian cell systems, and some have advanced to clinical studies (Huang et al., 2024). Several studies have shed light on the potential of tyrosine kinase inhibitors for TB treatment. For example, therapeutic doses of imatinib have proven effective in enhancing phagosome acidification and maturation, leading to a reduction in intracellular Mtb survival, both *in vitro* and in infected mice (Bruns et al., 2012; Napier et al., 2011). Another HDT approach involves metformin, a medication commonly prescribed for type 2 diabetes, which activates AMP-activated protein kinase. Metformin not only reduces the bacterial burden but also improves lung pathology in both mice and humans by enhancing autophagy and increasing the production of reactive oxygen species (Singhal et al., 2014). An emerging avenue focuses on inhibiting Mtb’s intracellular acquisition of fatty acids. Tetrahydrolipstatin (THL), which is recognized as an inhibitor of pancreatic lipase and marketed as a weight loss aid, shows promise as an anti-Mtb compound. Although the precise mechanism of its action is only partially understood, it is believed that THL depletes essential lipids from the intracellular environment, which are critical for Mtb’s survival (Parker et al., 2009).

The *trans*-δ-viniferin scaffold exerted the strongest anti-infective effect in our assay. Members of this family are phytoalexins found in various plants, including grapevines (*Vitis vinifera* L.), and they act as defense compounds produced by plants in response to stress, infection, or other environmental challenges, including microbial pathogens (Langcake & Pryce, 1976). Phenolic compounds such as stilbenes and the *trans*-δ-viniferins presented here have many reported biological activities against bacteria (Langcake & Pryce, 1976; Righi et al., 2020; Vestergaard & Ingmer, 2019), but also diverse positive effects on mammalian cells (Li et al., 2016; Qureshi et al., 2012; Ren et al., 2018; Rimando et al., 2005).

*Trans*-δ-viniferins belong to the family of polyphenols, which are known to non-selectively precipitate proteins (Siebert et al., 1996; Soares et al., 2012) and interfere with membrane structure (Borisova et al., 2019; Karonen, 2022). In previous work, urolithins and ellagitannins from plant extracts were already described as active in an amoebae-Mm infection model (E. A. Diop et al., 2018; E. H. A. Diop et al., 2019). We speculate that virulence factors of Mm, such as the ESX-1 effectors EsxA and EsxB (Gröschel et al., 2016), might be precipitated inside the MCV or the waxy outer membrane might be compromised during infection.

In a recent study, investigating essential requirements for infection of Mm in Dd via Transposon sequencing (Tn-Seq), we highlighted genes and pathways that might act as targets for antivirulence compounds (Lefrançois et al., 2024). Members of lipid metabolism are prominent candidates, such as genes required for phthiocerol dimycocerosates (PDIMs) synthesis (Quigley et al., 2017) or the *Mce* operons, which are associated with lipid transport (Klepp et al., 2022).

The host membrane might also be altered in a way that makes it less sensitive to the bacteria membrane damaging machinery, for example by interfering with cholesterol in the MCV membrane. Cholesterol has been discussed to play a role in membrane damage, bacterial entry and also during infection of other intracellular, vacuolar pathogens (De Jonge et al., 2007; Gatfield & Pieters, 2000; Howe & Heinzen, 2006). In addition, in Dd a knock-out of vacuolins, the functional homologues of the mammalian flotillins which organize membrane microdomains rich in sterols, lead to intracellular growth attenuation of Mm (Bosmani et al., 2021).

Since we did not observe a striking shift in IC_50_ upon manipulating the shape of the *trans*-δ-viniferins **17** and **19** via planarisation (benzofuran derivatives) and *cis*-isomerisation of the double bond, we speculate that the mechanism of action is unlikely to follow the conventional pattern of selective binding to a catalytic pocket on a protein but may indeed rely on interference with cellular membranes, as discussed above.

However, *trans*-δ-viniferins have been reported as Wnt modulators, which in turn is hypothesized to be a target for pharmaceutical autophagy induction (Huber, Koval, et al., 2022). Interestingly, autophagy is also discussed as a central process of cell autonomous defence against Mtb and Mm (Aylan et al., 2023; Golovkine et al., 2023).

Since *trans*-δ-viniferins show activities in many biological systems, this class of compounds might be described as pan assay interference compounds (PAINs) (Baell, 2016). PAINs are compounds that in principle can be developed into drugs, but due to their promiscuous target interactions in a less systematic manner and thus at a higher cost. For example, catecholamine drugs contain catechols, which are considered a classical feature of a PAIN (Baell, 2016). Other structural features of PAINs, such as a flexible three-dimensional shape and reactive side groups are present in *trans*-δ-viniferins. Nevertheless, we observed striking specific activity on intracellular Mm for **17**, **62**, **63** and **64**. In addition, current developments in pharmaceutical research try to embrace drugs with multiple targets, facilitated by advancing techniques to decipher complicated mechanisms of action, possibly allowing to progress putative PAINs more systematically, in the future.

A few structures were also found to enhance Mm growth during infection, but not in broth. During infection, **13, 14, 50, 51, 54** resulted in a normalized residual growth value greater than 1.5 (Fig. 1C, Table S1). Compounds that promote growth during infection, so-called pro-infectives may also be of interest. Latent TB is notoriously difficult to treat because antibiotics are more effective against bacteria in a growth phase. Mtb that enters the dormant phase can remain in the lung tissue for years and trigger a relapse to active TB (Bagcchi, 2023). In this context, compounds that can switch mycobacteria from a dormant to a growing state in a controlled manner might allow hitting these bacilli with classical therapeutic antibiotics, and may thus be of particular interest (Seidi & Jahanban-Esfahlan, 2013). On the other hand, such compounds might act by disabling cell-autonomous defence and thus might enhance our understanding of innate immune mechanisms. In addition, it is not clear how well the Dd-Mm platform reflects the dormant state of TB. Therefore, we wanted to focus on compounds that limit Mm growth and did not explore pro-infective compounds further in this study.

In summary, we emphasize the value of using nature inspired methods to derivatize simple scaffolds to bioactive compounds. Additionally, we demonstrated the use and power of the Dd-Mm infection system for drug discovery.

## ACKNOWLEDGEMENTS

We acknowledge Dr. Richard Fish from the Zebrafish Core Facility, Department of Genetic Medicine and Development, Faculty of Medicine, CMU, Geneva, Switzerland, for developing the zebrafish infection assay with us and performing initial experiments. We thank Emilie Michellod for the generation of the enzymatic secretome of *Botrytis cinerea* used on the biotransformation reactions. Furthermore, we thank Camille-Léa Dubuis for initial experiments with the library. We thank the Swiss National Science Foundation for providing financial support for this project, which aims to improve natural products chemical biodiversity for drug discovery by fungal secretome assisted biotransformation (grant 205321_182438/1 EFQ and KG). This work has also been supported by grants awarded to TS: A SystemsX grant (HostPathX), an SNF Sinergia grant (CRSII5_189921) and two SNF grants (310030_169386, 310030_188813).

## Abbreviation list

Dd: *Dictyostelium discoideum*
Mm: *Mycobacterium marinum*
Mtb: *Mycobacterium tuberculosis*
TB: Tuberculosis
SAR: Structure-activity relationship
NP: Natural product
WHO: World Health Organization
MDR: Multi-Drug-Resistant
IC_50_: Half maximal inhibitory concentration
DRC: Dose-response curve
AUC: Area under the curve
VS: Vehicle control
ECD: Electronic circular dichroism (ECD)
NMR: Nuclear magnetic resonance
HRMS: High-resolution mass spectrometry
HDT: Host-directed therapy
MCV: Mycobacteria-Containing Vacuole

## SUPPLEMENTARY MATERIAL

**Table S1. Activity data from single-dose assays and dose-response curves, compound references and RFU calibration data**

The file contains four sheets, with the first sheet “Legend” presenting a legend and detailed explanation of the remaining three sheets. Briefly, sheet two “Compounds” contains an overview over the library compounds and derivatives, their names, ids, properties, activities from screening at single dose experiments at 20µM and summary data from dose-response curves of selected compounds. Sheet three “Antibiotics” contains summary data from dose-response curves of benchmarked antibiotics in Fig. 1. Sheet four “Cell-RFU calibration” contains a calibration curve, demonstrating the linear relationship between random fluorescence units and amoeba cell density.

**Fig. S1.**
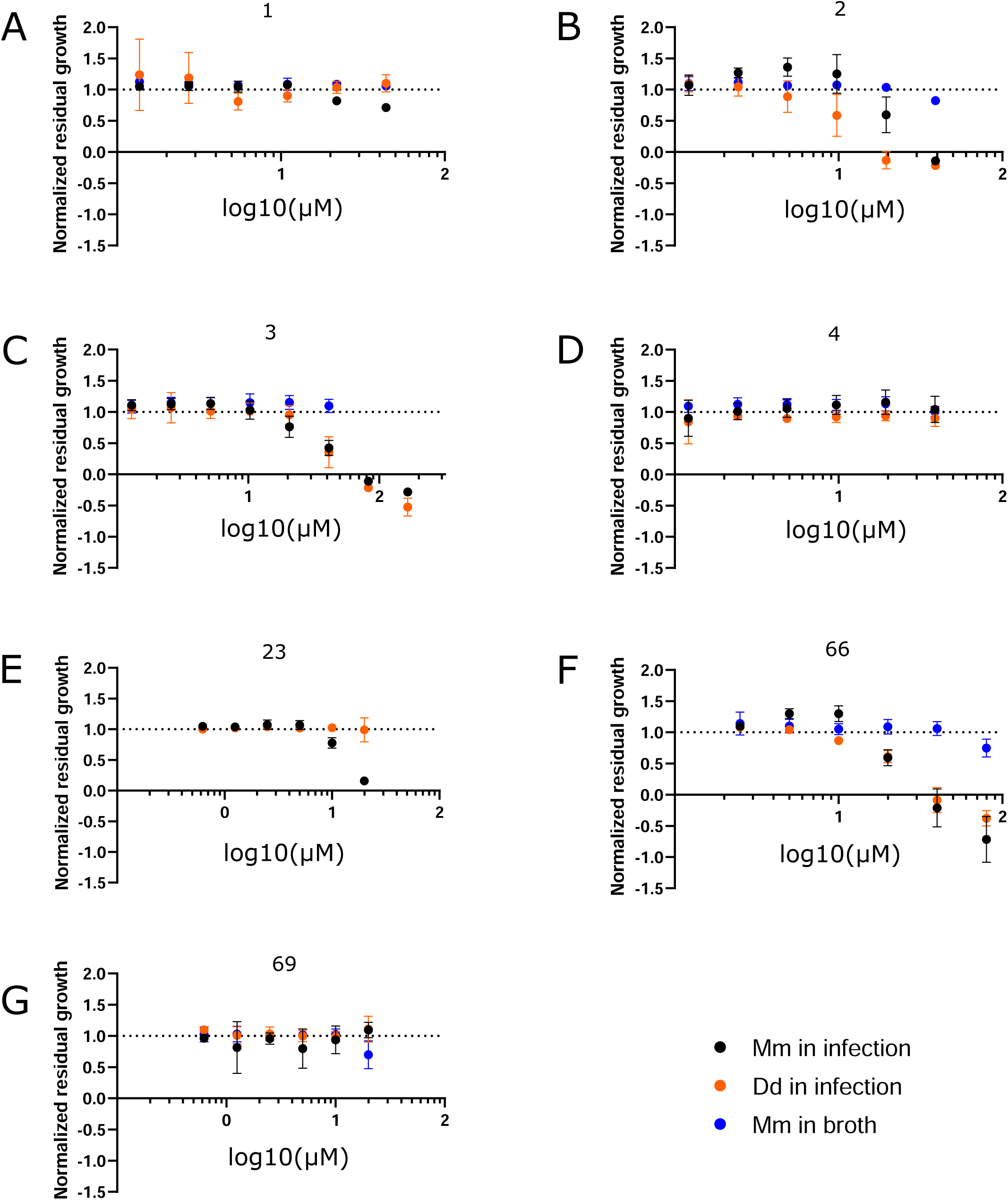
Stilbenes and *trans*-δ-viniferins derivatives structure activity relationship. Depicted are dose-response curves. **A-D** shows stilbene monomers, resveratrol (**1**), pterostilbene (**2**), pinostilbene (**3**) and isorhapontigenin (**4**), respectively. **B** shows Mm in infection and Dd in infection for 23, a hit compound from the initial library screen. F and G show benzofuran derivatives **66** and **69** of **11** and **21**, respectively. Black points show Mm in infection, orange points Dd in infection and blue points Mm in broth; Log10 of the concentration in µM is shown on the x-axis. A dashed line at y = 1 represents the normalized residual growth of the vehicle control. Depicted are means of at least three biological replicates and the respective standard deviations.

**Fig. S2.**
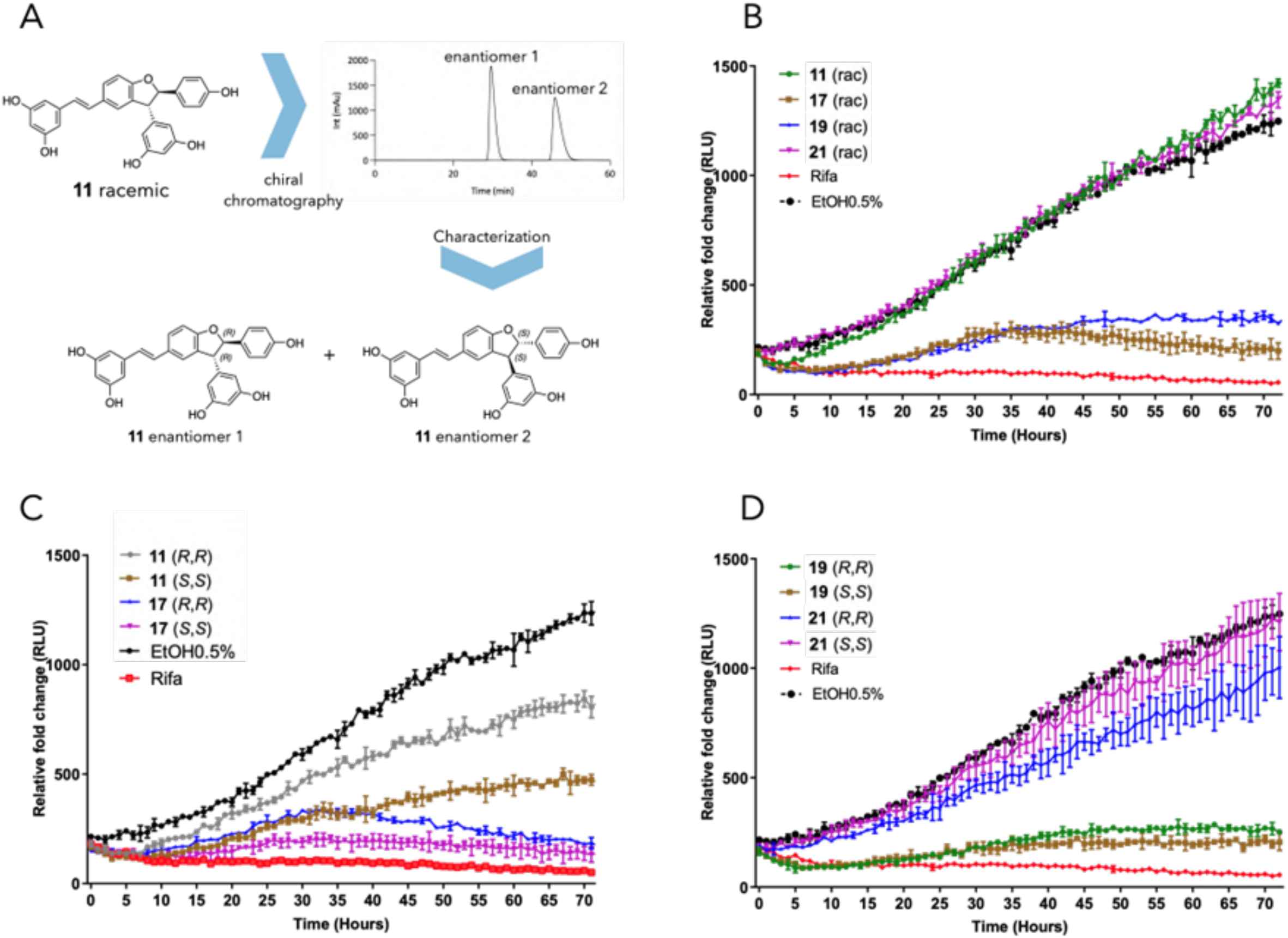
Enantiomer separation and activity. Panel **A** shows the generic approach to isolate the enantiomers of the *trans*-δ-viniferin, exemplified by the compound **11**. **B** shows Mm in infection growth curves of racemic mixtures, **C** Mm in infection growth curves of the enantiomers of **19** and **21**, **D** Mm in infection growth curves of the enantiomers of **11** and **17**.

**Fig. S3:**
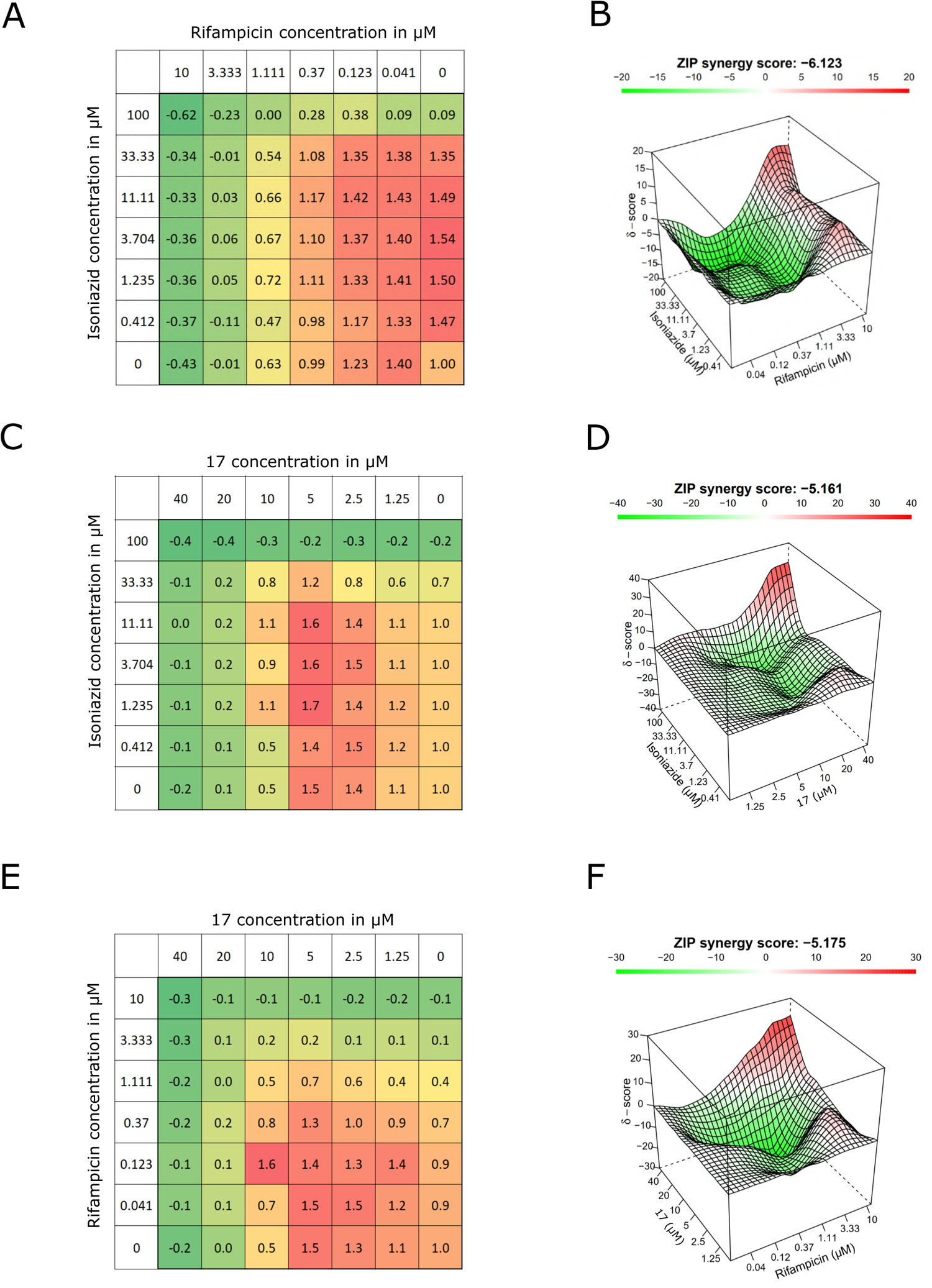
Checkerboard assays of rifampicin, isoniazid and 17. Synergy between **17** and the antibiotics isoniazid and rifampicin was investigated in checkerboard assays and subsequent analysis with synergyfinder using the ZIP synergy score. Panels **A**, **C** and **E** show normalized residual growth for every combination of concentrations for isoniazid and rifampicin, **17** and isoniazid, and **17** and rifampicin respectively. Green codes for low residual growth and red codes for high residual growth. Panels **B**, **D** and **F** show the computed ZIP synergy score and the concentration of the respective compound. Red codes for synergy, green codes for antagonism.

